# MosaicTR: tandem repeat somatic instability quantification from long-read sequencing

**DOI:** 10.64898/2026.03.16.712141

**Authors:** Junsoo Kim

## Abstract

**Summary:** Somatic instability of tandem repeats modifies disease onset and progression in repeat expansion disorders and serves as a biomarker for mismatch repair deficiency in cancer. MosaicTR quantifies per-locus somatic instability from haplotype-tagged long-read sequencing data without the read-length and PCR stutter limitations of short-read approaches. A motif-unit-weighted metric reduces platform-specific sequencing noise on both PacBio HiFi and Oxford Nanopore data, and pairwise comparison modes support detection of tissue-specific or longitudinal instability changes.

**Availability:** MosaicTR is freely available at https://github.com/junsoopablo/mosaictr under the MIT license.

## 1 Introduction

Somatic instability of tandem repeats (TRs) influences disease phenotype in repeat expansion disorders and cancer. In Huntington disease, somatic expansion of the CAG repeat in *HTT* correlates with disease onset and progression, and mismatch repair genes, particularly *MSH3*, have been identified as genetic modifiers of this process [1–3]. At the genome-wide scale, microsatellite instability (MSI) serves as a biomarker for mismatch repair deficiency and predicts response to immune checkpoint blockade in cancer [4]. Somatic expansion rates vary across tissues and accumulate over time [5], so characterizing instability requires profiling at the resolution of individual loci. Because repeat expansion disorders are typically heterozygous [6], distinguishing the expanded from the normal allele further demands haplotype-level analysis. Large-scale efforts such as the Somatic Mosaicism across Human Tissues (SMaHT) consortium [7] are now generating matched long-read sequencing data across multiple tissues from the same individuals, creating a need for tools that quantify instability at individual loci with haplotype resolution.

Several tools address components of this problem. MSI classifiers such as MSIsensor [8] and MANTIS [9] provide binary tumor-level status from short reads. prancSTR [10], search-STR [11], and MSMuTect [12] measure per-locus instability from short-read data, but are limited to repeats within read length. Long-read genotypers such as TRGT [13], LongTR [14], STRkit [15], and Straglr [16] provide accurate allele sizing across large expansions, and Owl [17] extends this to per-locus instability quantification from PacBio HiFi data using coefficient of variation. However, no existing tool combines motif-unit-aware noise separation, multi-platform support, and multi-tissue comparison for haplotype-resolved instability quantification.

MosaicTR operates on HP-tagged BAM files, a standard output of HiFi and nanopore variant-calling pipelines, and computes per-haplotype instability without additional phasing at each locus. Its primary metric, the Haplotype Instability Index (HII), measures intraallelic repeat length variation normalized by motif length, with motif-unit-aware weighting that attenuates platform-specific sequencing noise on both PacBio HiFi and Oxford Nanopore (ONT) data. For multi-tissue and longitudinal studies, pairwise comparison and matrix modes detect tissue-specific or passage-dependent instability changes.

## 2 Methods

### Read extraction and allele sizing

MosaicTR takes an HP-tagged BAM file and a four-column BED loci catalog as input. For each locus, reads overlapping the repeat region with ≥50 bp flanking sequence on both sides are extracted (mapping quality ≥5, up to 200 reads per locus). Allele sizes are computed from CIGAR strings as the sum of alignment operations spanning the repeat interval, without sequence-level realignment. Reads are grouped by HP tag. MapQ-weighted medians per haplotype, with conditional motif-unit rounding, yield diploid allele sizes. A concordance-based decision function determines zygosity. When HP tags are unavailable, MosaicTR falls back to gap-split analysis, which uses a bimodality test to separate reads into pseudohaplotypes, or to pooled analysis for unimodal distributions.

### Haplotype instability index

From the per-haplotype allele size distributions, instability is quantified using the Haplotype Instability Index:

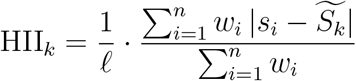

where *S*_*k*_ = {*s*_1_, …, *s*_*n*_} are the allele sizes assigned to haplotype 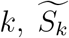 is the haplotype median, *𝓁* is the motif length, and *w*_*i*_ is a per-read weight based on motif-unit consistency: deviations that are integer multiples of *𝓁* receive full weight (*w*_*i*_ = 1), while sub-motif deviations are down-weighted (*w*_*i*_ = 0.1). This weighting reflects the error profile of PacBio HiFi sequencing, in which 92% of residual errors are sub-motif homopolymer indels [18], whereas somatic TR expansion and contraction occur predominantly in whole motif units [5]. A stable allele yields HII near zero. Somatic mosaicism elevates HII in proportion to repeat length heterogeneity within the haplotype. An auxiliary Inter-haplotype Asymmetry Score (IAS = |HII_1_ −HII_2_ |*/* max(HII_1_, HII_2_)) is also reported, ranging from 0 (symmetric instability) to 1 (instability confined to one haplotype). MAD-based outlier trimming removes misaligned or chimeric reads prior to metric computation.

### Oxford Nanopore noise model

Oxford Nanopore reads exhibit per-read random length jitter, predominantly ± 1 bp indels, that differs in character from the sub-motif homopolymer indels predominant on PacBio HiFi [19]. For repeat motifs ≥3 bp, this 1 bp jitter is sub-motif and receives reduced weight (*w* = 0.1). For dinucleotide repeats, however, a 2 bp ONT error equals one full motif unit and receives full weight, which elevates baseline noise. MosaicTR auto-detects the sequencing platform from the BAM header and applies motif-length-aware noise thresholds for ONT data: 2.0 for dinucleotides, decreasing to 0.5 for motifs ≥7 bp.

### Multi-sample instability comparison

For longitudinal or multi-tissue studies, a pairwise comparison mode identifies loci with instability changes between two samples (e.g., tumor–normal or early–late passage). A matrix mode computes all pairwise ΔHII values across an arbitrary number of samples.

### Simulation design

Synthetic read distributions were generated to validate HII recovery and classification performance. Because somatic expansions are more frequent than contractions at disease loci [1, 5], an expansion-biased model was used: for each simulated locus, read-level allele sizes were drawn as *s*_*i*_ = *m* + *X*_*i*_ + *ϵ*_*i*_, where *m* is the inherited allele size, *X*_*i*_ ~ Exponential(*λ*) models somatic expansion, and *ϵ*_*i*_ ~ 𝒩 (0, 0.5 bp) models measurement noise. The exponential scale *λ* was calibrated so that the expected absolute deviation equals the target HII × motif length. Stable loci were simulated with Gaussian noise only (*σ* = 0.5 bp). Dose-response analysis used target HII × values from 0 to 20 at 30× coverage with a 3 bp motif; ROC analysis used 500 stable and 500 unstable loci; coverage sweep varied per-haplotype depth from 5× to 80×.

### Datasets

Genome-wide noise characterization used HG002 PacBio HiFi Revio data (48 ×; Genome in a Bottle Consortium, NIST) aligned to GRCh38, with 108,584 tandem repeat loci from the Adotto v1.2 catalog [13]. Disease carrier validation used eight PureTarget reference standards (Coriell Institute) for Huntington disease (*HTT*), Fragile X syndrome (*FMR1*), SCA1 (*ATXN1*), SCA3 (*ATXN3*), myotonic dystrophy type 1 (*DMPK*), Friedreich ataxia (*FXN*), and FTD/ALS (*C9ORF72*), sequenced on PacBio HiFi. Oxford Nanopore carrier detection used 100 samples from the 1000 Genomes ONT dataset (R9.4.1 chemistry) [20], with HP tagging via LongPhase [21] using NYGC 30× Illumina phased VCFs as a SNP scaffold. An additional HTT carrier (HG02275) was identified from the 1000 Genomes ONT Vienna dataset and haplotagged with whatshap [22]. Longitudinal passage analysis used the HG008 pancreatic cancer cell line (normal tissue through passage 41; GIAB).

### Implementation

MosaicTR is implemented in Python and requires only pysam and NumPy as core dependencies. It provides a command-line interface with subcommands for genotyping, instability quantification, multi-sample comparison, and optional VCF output. Output is a 13-column tab-delimited file reporting per-haplotype allele sizes, HII, IAS, analysis path (HP-tagged, gap-split, or pooled), and read counts.

## 3 Results

The Haplotype Instability Index (HII) was validated on synthetic read distributions with known instability levels (Supplementary Figure S1). Measured HII scaled linearly with simulated instability (*R*^2^ = 1.000), with ~ 30% attenuation because outlier trimming and motif-unit down-weighting reduce the contribution of extreme or sub-motif deviations. For binary stable/unstable classification, ROC analysis gave an AUC of 0.975 (99.3% sensitivity, 100% specificity) at a threshold of HII = 0.45. Detection of 80% of unstable loci required 15× per-haplotype depth.

The noise baseline was characterized at 108,584 tandem repeat loci in the healthy reference sample HG002 (Genome in a Bottle) on PacBio HiFi. The median HII was 0.004, and 99% of loci fell below the 0.45 threshold, which corresponds to the 3*σ* upper bound of this distribution. Motif-unit weighting reduced the false positive rate from 10.2% to 0.9% at these loci (Supplementary Figure S4a). HP-tagged analysis further reduced false positive instability calls by 70% at heterozygous loci, where allele length differences otherwise inflate apparent instability (Supplementary Figure S3). At 10,503 coordinate-matched loci, MosaicTR’s noise distribution was narrower than that of Owl [17], which uses unweighted coefficient of variation (Supplementary Figure S4b). On ONT, the motif-unit weighting adapted to platform-specific noise: ONT reads exhibit per-read ± 1 bp length jitter [19], which is sub-motif for repeats ≥3 bp and accordingly receives reduced weight. This noise scaled inversely with motif length (mean HII: dinucleotide 0.62, trinucleotide 0.39, pentanucleotide 0.20 across 2,625 loci at non-carrier samples in the 1000 Genomes dataset), and at ATXN10 loci (ATTCT, 5 bp motif) normal allele HII on ONT was 0.03–0.04, approaching HiFi levels (Supplementary Figure S2c).

With this noise baseline established, MosaicTR was applied to disease carrier detection. Nine of twelve tested samples showed detectable instability across five disorders (Huntington disease, SCA10, Fragile X, SCA3, SCA1) on PacBio HiFi and Oxford Nanopore platforms; three PureTarget samples had expanded allele dropout (Figure 1C; Supplementary Table S1). Three ATXN10 carriers were identified from 100 samples in the 1000 Genomes ONT dataset (Figure 1B). Among these, expanded allele HII scaled with expansion size: HG02345 (321 repeat units) had HII = 0.6, HG02252 (944) had 4.3, and HG01122 (1,041) had 31.0. This gradient matches somatic mosaicism observed by Southern blot in SCA10 [23] and the length-dependent instability reported in Huntington disease [5]. The read-level allele size distribution of HG01122 on the expanded haplotype followed an exponential distribution (Kolmogorov–Smirnov test, *p* = 0.67; Supplementary Figure S2a), consistent with the expansion-biased simulation model. All carrier HII values exceeded the detection threshold, with signal-to-noise ratios of 115- to 5,964-fold on HiFi and 13- to 17-fold on ONT relative to each platform’s baseline. Haplotype resolution distinguished heterozygous carriers, where instability is confined to one allele (IAS ≈ 1), from a bilateral expansion in HG02252 where both alleles were expanded and somatically unstable (IAS = 0.12), a distinction inaccessible from pooled read data.

**Figure 1.**
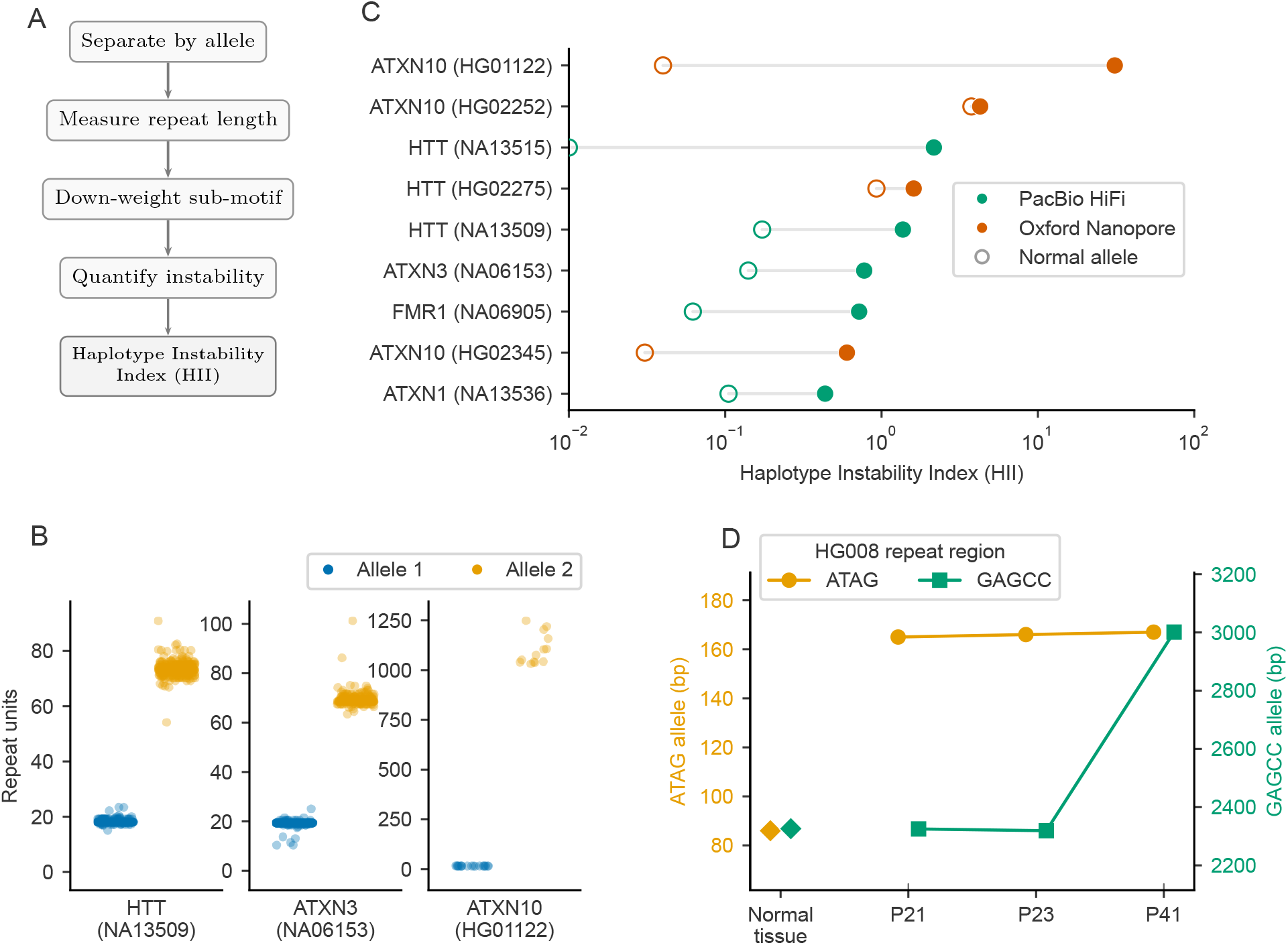
MosaicTR overview and validation. (A) Algorithm overview: HP-tagged reads are separated by allele, repeat lengths are measured from CIGAR strings, sub-motif deviations are down-weighted to suppress sequencing noise, and per-haplotype instability (HII) is computed. (B) Per-read allele sizes for three disease carriers on PacBio HiFi (HTT, ATXN3) and Oxford Nanopore (ATXN10). Blue: allele 1; orange: allele 2. Each dot represents a single read. (C) Haplotype Instability Index (HII) across nine expansion carriers spanning five repeat disorders. Open circles: normal allele; filled circles: expanded allele. Green: PacBio HiFi; orange: Oxford Nanopore. HII spans three orders of magnitude, with expanded alleles consistently elevated above normal alleles. (D) Allele size trajectories at two repeat loci across HG008 pancreatic cancer cell line passages (normal tissue through passage 41), showing passage-dependent size changes.

Beyond static carrier detection, MosaicTR’s comparison mode was used to track longitudinal instability changes in the HG008 pancreatic ductal adenocarcinoma cell line (Genome in a Bottle; normal tissue through passage 41). Several loci showed progressive expansion: ATAG at chr22:11.6M grew from 86 to 167 bp, and GAGCC at chr21:8.4M gained +675 bp between passages 23 and 41 (Figure 1D). Of 1,049 drifted loci (ΔHII ≥0.5), 90% were dinucleotide repeats, consistent with replication slippage at short motifs [24]. No mononucleotide instability was observed, as expected for a microsatellite-stable cell line in which intact mismatch repair suppresses mononucleotide repeat slippage.

## 4 Discussion

We presented MosaicTR, a tool that quantifies somatic tandem repeat instability from long-read sequencing data by combining haplotype-aware analysis, motif-unit-weighted instability scoring, and inter-haplotype asymmetry measurement. Existing short-read instability tools such as prancSTR [10] are limited to expansions within read length (~ 150 bp) and subject to PCR stutter artifacts at trinucleotide repeats [25]. Long-read genotypers such as TRGT, LongTR, and Straglr [13, 14, 16] have established accurate allele sizing that spans disease-relevant expansions, and Owl [17] extended this to per-locus instability quantification using coefficient of variation. MosaicTR builds on these advances by exploiting the error profile of long-read platforms: on PacBio HiFi, 92% of residual errors are sub-motif homopolymer indels [18], whereas somatic repeat length changes occur in whole motif units. Weighting reads by this motif-unit consistency separates sequencing noise from biological signal, reducing the false positive rate to 0.9% at 108,584 loci (Supplementary Figure S4).

Several limitations should be noted. MosaicTR requires HP-tagged BAMs, which are standard output from PacBio DeepVariant pipelines but may need an additional phasing step on ONT and other platforms. Dinucleotide repeats on ONT remain challenging because a 2 bp indel equals one full motif unit and cannot be down-weighted; the ONT results reported here used R9.4.1 chemistry [20], and the newer R10.4.1 chemistry [26] may reduce this noise. Other limitations include very large expansions (*>*10 kb) that may exceed read lengths, and VNTR alignment ambiguity that limits sizing precision for complex repeats.

Multi-tissue long-read datasets are becoming available through efforts such as SMaHT [7], which is generating matched tissue panels from hundreds of donors with both PacBio HiFi and ONT sequencing. MosaicTR’s pairwise comparison and matrix modes are designed to quantify per-locus instability differences across such tissue panels, enabling systematic characterization of tissue-specific and age-dependent somatic expansion patterns at tandem repeat loci genome-wide.

---

**Table 1:**
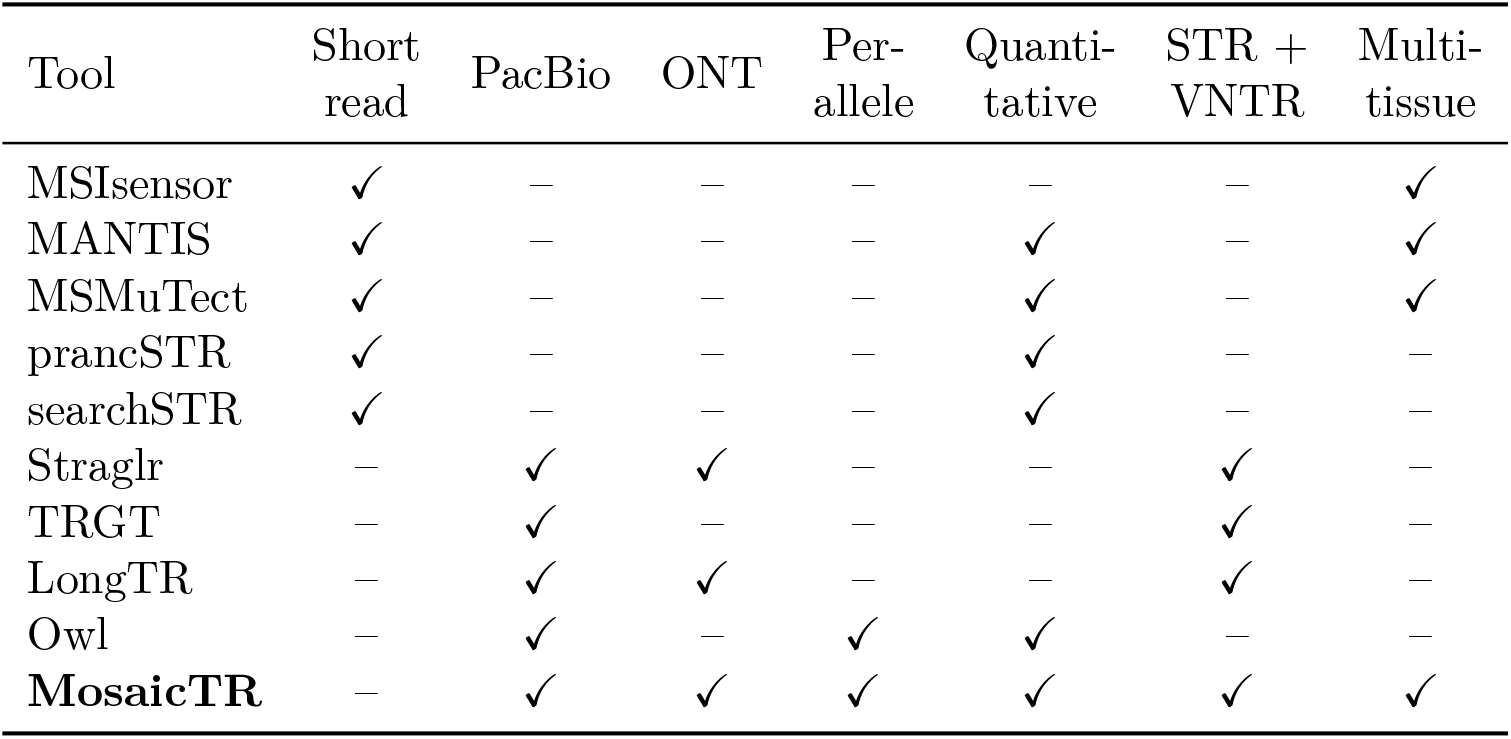
Comparison of tandem repeat instability tools. “Per-allele” indicates whether instability is resolved for each haplotype separately. “Quantitative” indicates a per-locus numeric instability score (as opposed to binary MSI classification). “Multi-tissue” indicates support for paired or multi-sample comparison (e.g., tumor–normal or cross-tissue).

| Tool | Short<br>read | PacBio | ONT | Per-allele | Quanti-<br>tative | STR +<br>VNTR | Multi-tissue |
| --- | --- | --- | --- | --- | --- | --- | --- |
| MSIsensor | ✓ | — | — | — | — | — | ✓ |
| MANTIS | ✓ | — | — | — | ✓ | — | ✓ |
| MSMuTect | ✓ | — | — | — | ✓ | — | ✓ |
| prancSTR | ✓ | — | — | — | ✓ | — | — |
| searchSTR | ✓ | — | — | — | ✓ | — | — |
| Straglr | — | ✓ | ✓ | — | — | ✓ | — |
| TRGT | — | ✓ | — | — | — | ✓ | — |
| LongTR | — | ✓ | ✓ | — | — | ✓ | — |
| Owl | — | ✓ | — | ✓ | ✓ | — | — |
| **MosaicTR** | — | ✓ | ✓ | ✓ | ✓ | ✓ | ✓ |

## References

[1] Meera Swami, Ashley E. Hendricks, Tammy Gillis, et al. Somatic expansion of the Huntington’s disease CAG repeat in the brain is associated with an earlier age of disease onset. Hum. Mol. Genet., 18:3039–3047, 2009. doi: 10.1093/hmg/ddp242.

[2] Marc Ciosi, Alison Maxwell, Sarah A. Cumming, et al. A genetic association study of glutamine-encoding DNA sequence structures, somatic CAG expansion, and DNA repair gene variants, with Huntington disease clinical outcomes. EBioMedicine, 48: 568–580, 2019. doi: 10.1016/j.ebiom.2019.09.020.

[3] GeM-HD Consortium. CAG repeat not polyglutamine length determines timing of Huntington’s disease onset. Cell, 178:887–900, 2019. doi: 10.1016/j.cell.2019.06.036.

[4] Dung T. Le, Jennifer N. Uram, Hao Wang, et al. PD-1 blockade in tumors with mismatch-repair deficiency. N. Engl. J. Med., 372:2509–2520, 2015. doi: 10.1056/NEJMoa1500596.

[5] Lesley Kennedy, Erin Evans, Chiun-Min Chen, et al. Dramatic tissue-specific mutation length increases are an early molecular event in Huntington disease pathogenesis. Hum. Mol. Genet., 12:3359–3367, 2003. doi: 10.1093/hmg/ddg352.

[6] Christel Depienne and Jean-Louis Mandel. 30 years of repeat expansion disorders: What have we learned and what are the remaining challenges? Am. J. Hum. Genet., 108:764–785, 2021. doi: 10.1016/j.ajhg.2021.03.011.

[7] SMaHT Network. The Somatic Mosaicism across Human Tissues Network. Nature, 2025. doi: 10.1038/s41586-025-09096-7.

[8] Beifang Niu, Kai Ye, Qunyuan Zhang, et al. MSIsensor: microsatellite instability detection using paired tumor–normal sequence data. Bioinformatics, 30:1015–1016, 2014. doi: 10.1093/bioinformatics/btt755.

[9] Esko A. Kautto, Russell Bonneville, Jharna Miya, et al. Performance evaluation for rapid detection of pan-cancer microsatellite instability with MANTIS. Oncotarget, 8: 7452–7463, 2017. doi: 10.18632/oncotarget.13918.

[10] Asmita Sehgal, Helyaneh Ziaei Jam, and Melissa Gymrek. prancSTR: genome-wide analysis of short tandem repeat mosaicism from short reads. bioRxiv, 2024. doi: 10.1101/2024.12.21.629930.

[11] Mir Faisal Aziz, Akdes Serin Dincer, et al. searchSTR: genome-wide profiling of short tandem repeat mosaicism. bioRxiv, 2025. doi: 10.1101/2025.01.13.632808.

[12] Yosef E. Maruvka, Kent W. Mouw, Monica Karber, et al. Analysis of somatic microsatellite indels identifies driver events in human tumors. Nat. Biotechnol., 35:951–959, 2017. doi: 10.1038/nbt.3966.

[13] Egor Dolzhenko, Viraj Deshpande, Felix Schlesinger, et al. Characterization and visualization of tandem repeats at genome scale. Nat. Biotechnol., 42:1606–1614, 2024. doi: 10.1038/s41587-023-02057-3.

[14] Helyaneh Ziaei Jam, Yang Li, Ross DeVore, et al. LongTR: genome-wide profiling of genetic variation at tandem repeats from long reads. Genome Biol., 25:176, 2024. doi: 10.1186/s13059-024-03319-2.

[15] Qian Liu, Xiangyu Tong, Disha K. Bhatt, et al. STRkit: a comprehensive toolkit for STR genotyping and analysis from long reads. Genome Biol., 25:200, 2024. doi: 10.1186/s13059-024-03345-0.

[16] Readman Chiu, Indhu-Shree Rajan-Babu, Jean Friedlander, et al. Straglr: discovering and genotyping tandem repeat expansions using whole genome long-read sequences. Genome Biol., 22:224, 2021. doi: 10.1186/s13059-021-02447-3.

[17] Zev Kronenberg, Khi Pin Chua, Mark J.P. Chaisson, et al. Hunting for microsatellite instability in long-read data with Owl. bioRxiv, 2026. doi: 10.64898/2026.03.08.708905. Preprint.

[18] Aaron M. Wenger, Paul Peluso, Richard J. Hall, et al. Accurate circular consensus long-read sequencing improves variant detection and assembly of a human genome. Nat. Biotechnol., 37:1155–1162, 2019. doi: 10.1038/s41587-019-0217-9.

[19] Clara Delahaye and Jacques Nicolas. Sequencing DNA with nanopores: troubles and biases. PLoS ONE, 16:e0257521, 2021. doi: 10.1371/journal.pone.0257521.

[20] Julia A. Gustafson, S. Blake Gibson, Nidhi Damaraju, et al. Nanopore sequencing of 1,000 genomes project samples. bioRxiv, 2024. doi: 10.1101/2024.03.05.583625.

[21] Yun-Lung Lin, Po-Chun Chang, Chien-Li Hsu, et al. LongPhase: an ultra-fast chromosome-scale phasing algorithm for the third-generation long read sequencing. Bioinformatics, 38:i203–i210, 2022. doi: 10.1093/bioinformatics/btac250.

[22] Marcel Martin, Patterson Murray, Shilpa Garg, et al. WhatsHap: fast and accurate read-based phasing. bioRxiv, 2023. doi: 10.1101/085050.

[23] Tohru Matsuura, Ping Fang, Xia Lin, et al. Somatic and germline instability of the ATTCT repeat in spinocerebellar ataxia type 10. Am. J. Hum. Genet., 74:1216–1224, 2004. doi: 10.1086/421526.

[24] Hans Ellegren. Microsatellites: simple sequences with complex evolution. Nat. Rev. Genet., 5:435–445, 2004. doi: 10.1038/nrg1348.

[25] Deepali Shinde, Yaling Lai, Fengzhu Sun, and Norman Arnheim. Taq DNA polymerase slippage mutation rates measured by PCR and quasi-likelihood analysis: (CA/GT)n and (A/T)n microsatellites. Nucleic Acids Res., 31:974–980, 2003. doi: 10.1093/nar/gkg178.

[26] Oxford Nanopore Technologies. R10.4.1 chemistry: improved accuracy for homopolymers and low-complexity sequences. Oxford Nanopore Technologies Technical Note, 2024. https://nanoporetech.com/accuracy.

