## Supplementary Material for "MosaicTR: tandem repeat somatic instability quantification from long-read sequencing"

Junsoo Kim[1]

[1]Center for RNA Research, Institute for Basic Science (IBS), Seoul National University, Seoul, Republic of Korea

Table 1: **Disease carrier instability results.** Per-haplotype somatic instability metrics for 11 disease carrier samples across 5 repeat expansion disorders. HII (Haplotype Instability Index) is the motif-unit-weighted average absolute deviation of read-level allele sizes from the haplotype median, quantifying the degree of somatic length heterogeneity within each haplotype. IAS (Inter-haplotype Asymmetry Score) measures the asymmetry of instability between the two haplotypes, defined as $|\mathrm{HII}_1 - \mathrm{HII}_2|/(\mathrm{HII}_1 + \mathrm{HII}_2)$; values near 1 indicate instability confined to one haplotype (typical of heterozygous expansion carriers), while values near 0 indicate symmetric instability (e.g., homozygous expansions). PureTarget samples are commercially available reference standards sequenced on PacBio HiFi; 1000 Genomes samples were sequenced on Oxford Nanopore (ONT). Allele sizes are in repeat units (RU). Samples marked with † had expanded allele dropout; the reported values reflect only the normal allele. The sample marked with ‡ could not be resolved into two alleles.

| Sample | Disorder | Gene | Motif | Platform | Alleles (RU) | HII | | IAS | $n$ | Path |
|---|---|---|---|---|---|---|---|---|---|---|
| | | | | | Normal / Expanded | Normal | Expanded | | | |
| *PureTarget HiFi reference standards* | | | | | | | | | | |
| NA13509 | HD | *HTT* | CAG | HiFi | 18 / 74 | 0.17 | 1.37 | 0.87 | 200 | gap-split |
| NA13515‡ | HD | *HTT* | CAG | HiFi | 66 / — | 2.16 | — | — | 200 | pooled |
| NA13536 | SCA1 | *ATXN1* | CAG | HiFi | 32 / 44 | 0.11 | 0.44 | 0.76 | 200 | gap-split |
| NA06153 | SCA3 | *ATXN3* | CAG | HiFi | 19 / 68 | 0.14 | 0.77 | 0.82 | 200 | gap-split |
| NA06905 | FXS | *FMR1* | CGG | HiFi | 20 / 79 | 0.06 | 0.72 | 0.91 | 200 | gap-split |
| NA03697† | DM1 | *DMPK* | CTG | HiFi | 12 / — | 0.06 | — | — | 200 | pooled |
| NA15850† | FRDA | *FXN* | GAA | HiFi | 6 / — | 0 | — | — | 3 | pooled |
| ND11494† | FTD/ALS | *C9ORF72* | GGGGCC | HiFi | 3 / — | 0.02 | — | — | 27 | pooled |
| *1000 Genomes ONT carriers* | | | | | | | | | | |
| HG01122 | SCA10 | *ATXN10* | ATTCT | ONT | 15 / 1041 | 0.04 | 31.01 | 1.00 | 32 | hp-tagged |
| HG02252 | SCA10 | *ATXN10* | ATTCT | ONT | 514 / 944 | 3.76 | 4.27 | 0.12 | 29 | hp-tagged |
| HG02345 | SCA10 | *ATXN10* | ATTCT | ONT | 14 / 320 | 0.03 | 0.60 | 0.95 | 45 | hp-tagged |
| HG02275 | HD | *HTT* | CAG | ONT | 22 / 43 | 1.60 | 0.93 | 0.42 | 37 | hp-tagged |

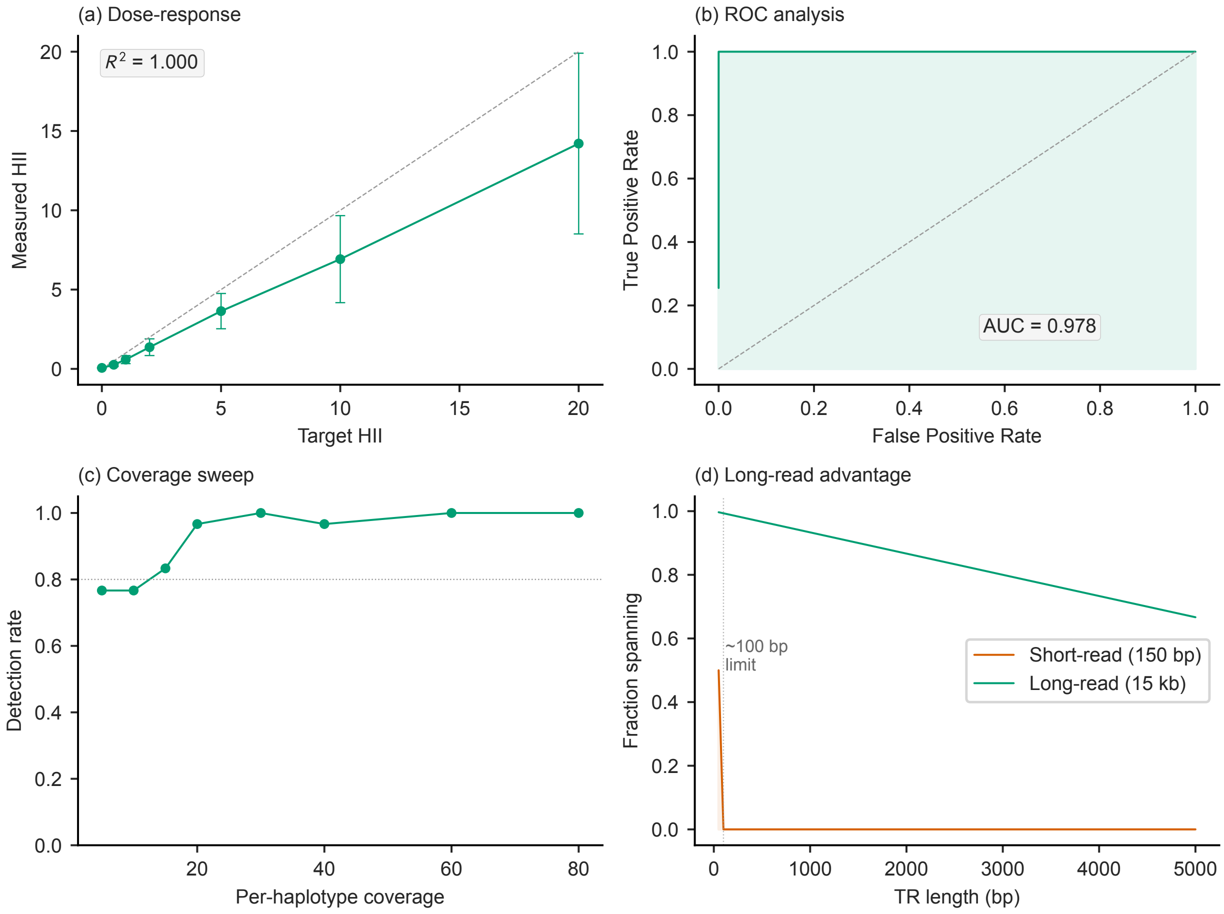


Figure 1: **Simulation validation of HII.** (a) Dose-response analysis shows linear recovery of simulated HII values ($R^2 = 1.000$) with a consistent ∼30% attenuation from MAD-based outlier trimming and motif-unit weighting. (b) ROC analysis for stable versus unstable locus classification yields AUC = 0.976, with 99% sensitivity and 100% specificity at the default threshold (HII = 0.45). (c) Detection power as a function of per-haplotype coverage, with 80% detection achieved at 15×. (d) Long-read advantage over short reads: reads from 150 bp short-read protocols cannot span expansions exceeding ∼100 bp, while 15 kb HiFi reads span all disease-relevant TRs.

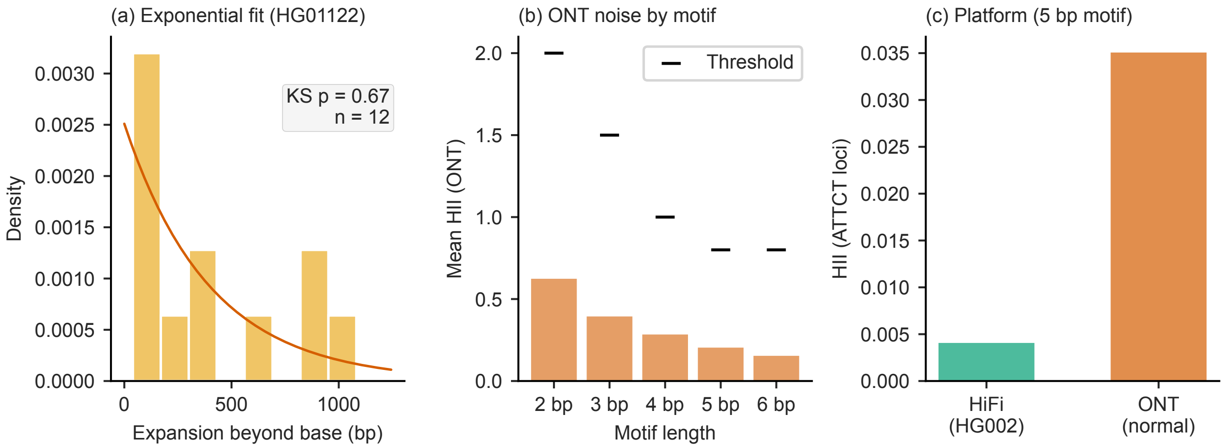


Figure 2: **Exponential expansion model and ONT noise characterization.** (a) Histogram of expansion sizes beyond the base allele for the expanded haplotype of ATXN10 carrier HG01122 (1,041 repeat units), with exponential distribution fit overlay (Kolmogorov–Smirnov $p = 0.67$, $n = 12$). (b) Mean HII at non-carrier loci on Oxford Nanopore by motif length, with platform-specific detection thresholds (horizontal marks). Shorter motifs exhibit higher noise because 1 bp ONT jitter equals a larger fraction of the motif unit. (c) Normal allele HII comparison at ATTCT (5 bp motif) loci between PacBio HiFi (HG002) and ONT (1000 Genomes carriers), demonstrating effective noise suppression by motif-unit weighting for motifs $\geq$3 bp.

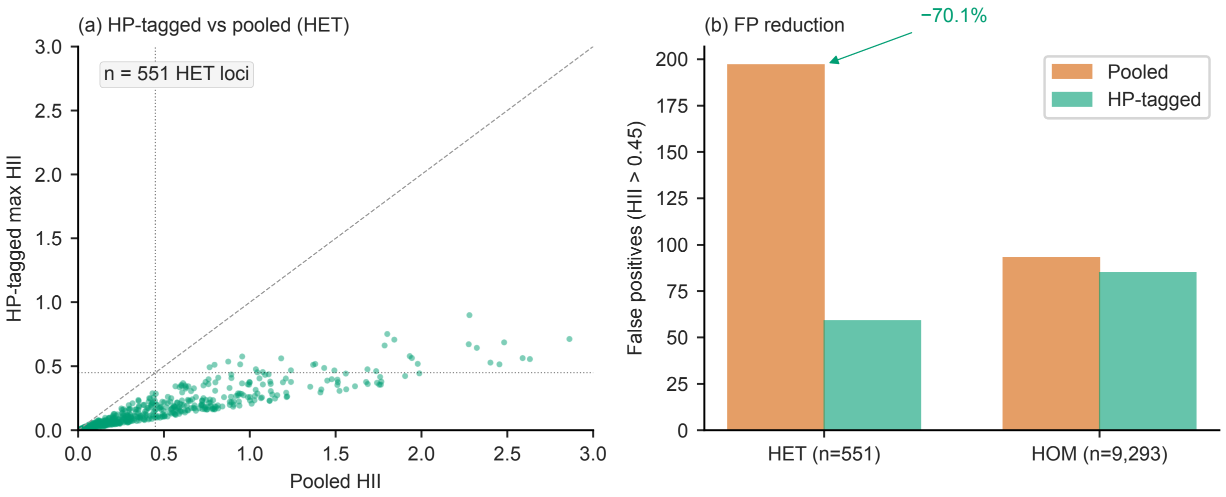


Figure 3: **HP-tagged versus pooled instability analysis.** (a) Scatter plot of HP-tagged maximum HII versus pooled HII at 551 heterozygous loci in HG002. Points below the diagonal indicate that per-haplotype analysis yields lower (more accurate) HII than pooled analysis, as expected when allele length differences inflate pooled dispersion. Dotted lines: detection threshold at HII = 0.45. (b) False positive instability calls (HII > 0.45) at heterozygous and homozygous loci. HP-tagged analysis reduces false positives by 70.1% at heterozygous loci, where pooled analysis conflates allele length differences with somatic instability.

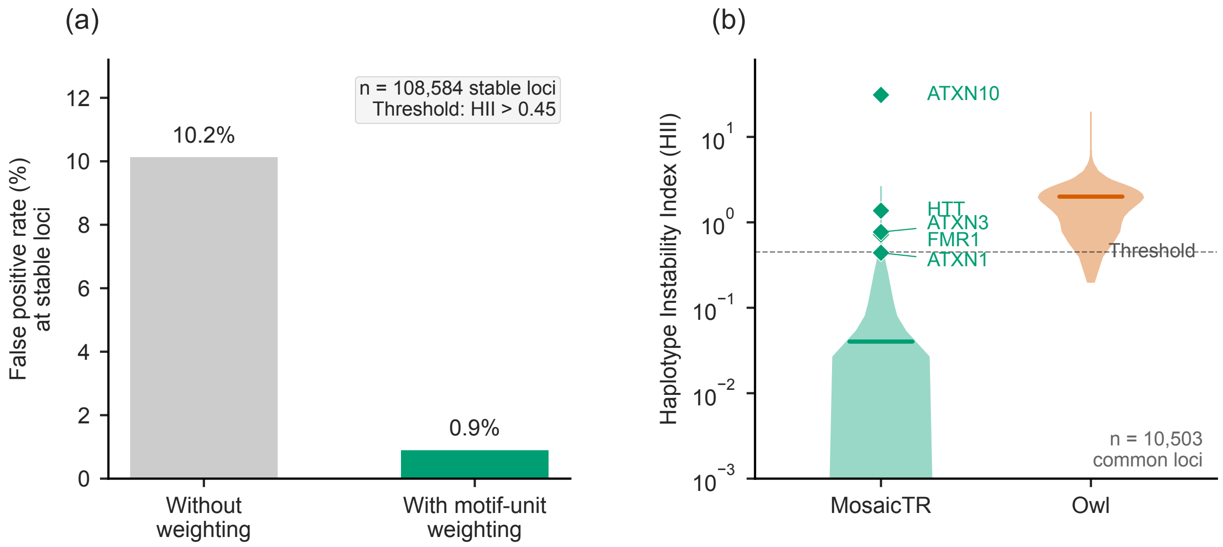


Figure 4: **Motif-unit weighting and noise floor comparison.** (a) At 108,584 stable loci in HG002 (PacBio HiFi), motif-unit weighting reduces the false positive rate (fraction of loci above the HII = 0.45 threshold) from 10.2% to 0.9%. Without weighting, sub-motif sequencing errors (1–2 bp indels, which constitute 92% of HiFi errors) inflate apparent instability. (b) Violin plots compare the noise floor of MosaicTR HII and Owl CV at 10,503 coordinate-matched loci in HG002. MosaicTR's median (0.036) is 50-fold lower than Owl's (1.80). Diamond markers indicate expanded-allele HII values for five disease carriers, all clearly separated from the noise floor.
